# SynLethDB 2.0: A web-based knowledge graph database on synthetic lethality for novel anticancer drug discovery

**DOI:** 10.1101/2021.12.28.474346

**Authors:** Jie Wang, Min Wu, Xuhui Huang, Li Wang, Sophia Zhang, Hui Liu, Jie Zheng

## Abstract

Two genes are synthetic lethal if mutations in both genes result in impaired cell viability, while mutation of either gene does not affect the cell survival. The potential usage of synthetic lethality (SL) in anticancer therapeutics has attracted many researchers to identify synthetic lethal gene pairs. To include newly identified SLs and more related knowledge, we present a new version of the SynLethDB database to facilitate the discovery of clinically relevant SLs. We extended the first version of SynLethDB database significantly by including new SLs identified through CRISPR screening, a knowledge graph about human SLs, and new web interface, etc. Over 16,000 new SLs and 26 types of other relationships have been added, encompassing relationships among 14,100 genes, 53 cancers, and 1,898 drugs, etc. Moreover, a brand-new web interface has been developed to include modules such as SL query by disease or compound, SL partner gene set enrichment analysis and knowledge graph browsing through a dynamic graph viewer. The data can be downloaded directly from the website or through the RESTful APIs. The database is accessible online at http://synlethdb.sist.shanghaitech.edu.cn/v2.

## Introduction

Synthetic lethality (SL), initially described in *Drosophila* as recessive lethality [9], is a type of gene-gene interaction such that the perturbation of both genes causes the loss of cell viability, while the perturbation of either gene alone will not affect the cell viability [31]. SL offers a strategy for cancer medicine by identifying new antibiotic or therapeutic targets [15, 3, 36]. By inhibiting the SL partner of a gene with cancer-specific alteration, we can kill cancer cells and spare normal cells, thereby reducing the side effect of the treatment [24, 23]. To discover SL gene pairs as a gold mine of cancer drug targets, researchers have applied various techniques, including chemical screening [16], RNAi screening [11, 1, 5, 29, 2], CRISPR screening [14, 39] and bioinformatics methods [22, 25, 41, 49].

The first version of SynLethDB released in 2016 contains 34,089 SL gene pairs and is the first comprehensive database of SLs [13]. It collects SL pairs for human and 4 model species, i.e., mouse, fruit fly, worm and yeast, from biochemical assays, public databases [38, 32], computational predictions [37] and text mining. In addition, it provides a statistical analysis module to evaluate the druggability and efficacy of SL pairs upon drug treatments by analyzing the large-scale drug sensitivity data. Recently, SynLethDB has been used as ground-truth SL data in various studies. For example, Liany et al. [26], Cai et al. [4] and Das et al. [7] used SynLethDB to train and test their computational SL prediction methods. Hu et al. [19] used SynLethDB to evaluate their method for *de novo* identification of synergistic optimal control nodes (OCNs) as candidate targets for combination therapy. Wang et al. [44] used the SLs in SynLethDB to investigate the link between SL interactions and drug sensitivity of cancer cells. Cui et al. used the SL data from SynLethDB in their web-based tool called siGCD [6] for analysis and visualization of the interactions among genes, cells and drugs associated with survival in human cancers.

Many CRISPR-based screening experiments have been conducted after 2015 and generated a large amount of data. Combinatorial CRISPR-based screening has been used to study genetic interactions, including the identification of SL interactions [48, 52, 39, 14, 46, 42]. Computational methods such as GEMINI [50] were proposed to identify SL gene pairs from these screening data. GEMINI is a variational Bayesian approach proposed to identify SLs from combinatorial CRISPR screens. Data driven method ISLE [25] searches in the lab-identified candidate SLs by tumor molecular profiles, patient clinical data, and gene phylogeny relations to find out the clinical SLs. These wet-lab experiments and computational methods provided further evidence for some existing SLs or discovered new SLs that had not been included in the first version of SynLethDB.

To discover SL-based anticancer drug targets and clinical SLs, it is highly desirable to consider the relationships among SLs, cancers and drugs. Several studies combined SLs with the information about cancers and cancer-drug interactions to discover cancer-specific SLs for new cancer therapies. SL-BioDP [8] provides an online tool based on a data-driven method to predict SL interactions by mining cancer’s genomic and chemical interactions. However, it only supports the prediction of SL partners of the 623 genes belonging to 10 hallmark cancer pathways and 18 types of cancers. SLKG [51] is a knowledge graph that contains 7 kinds of relationships among genes, cancers and drugs. Unlike SL-BioDP, SLKG collects SL pairs from literature and existing databases instead of by prediction. Moreover, SLKG is also used to identify the best repurposable drug candidates and drug combinations.

In addition to the relationships among SLs, drugs, and cancers, their various features are also useful for discovering SLs and anticancer therapy. Taking the features of genes as an example, the co-expression, gene ontology (GO) semantic similarity, and shared pathways between genes are commonly used features for predicting SLs [22, 7, 40, 27, 20]. In addition, several tools have been developed to curate these features, such as GO terms and pathways associated with specific genes, anatomies and symptoms of cancers, and the side effects and pharmacologic classes of drugs. For example, the Hetio package from Hetionet [17] provides a way to integrate different resources into a single data structure. We are motivated to use these tools to construct an integrative knowledge graph to better describe SL pairs.

In this paper, we present SynLethDB 2.0 to include newly discovered SLs and provide more related knowledge to help identify clinically relevant SLs (Figure 1). It is a significant expansion of the first version by adding 16,781 new SL gene pairs, and integrating a biomedical knowledge graph, including 10 kinds of biomedical entities other than gene and 26 kinds of relationships for drug discovery other than SL. The 37,341 entities and 1,405,652 relationships were used to create a knowledge graph and stored in a graph database. A user-friendly website interface with new functionalities for data browsing, visualization and analysis has also been developed for users to browse the data and knowledge graph in SynLethDB. For example, users can search SLs by a disease or a compound, perform pathway or GO term enrichment for SL partners of a gene, and inspect the connections between two genes in a interactive viewer.

**Fig. 1.**
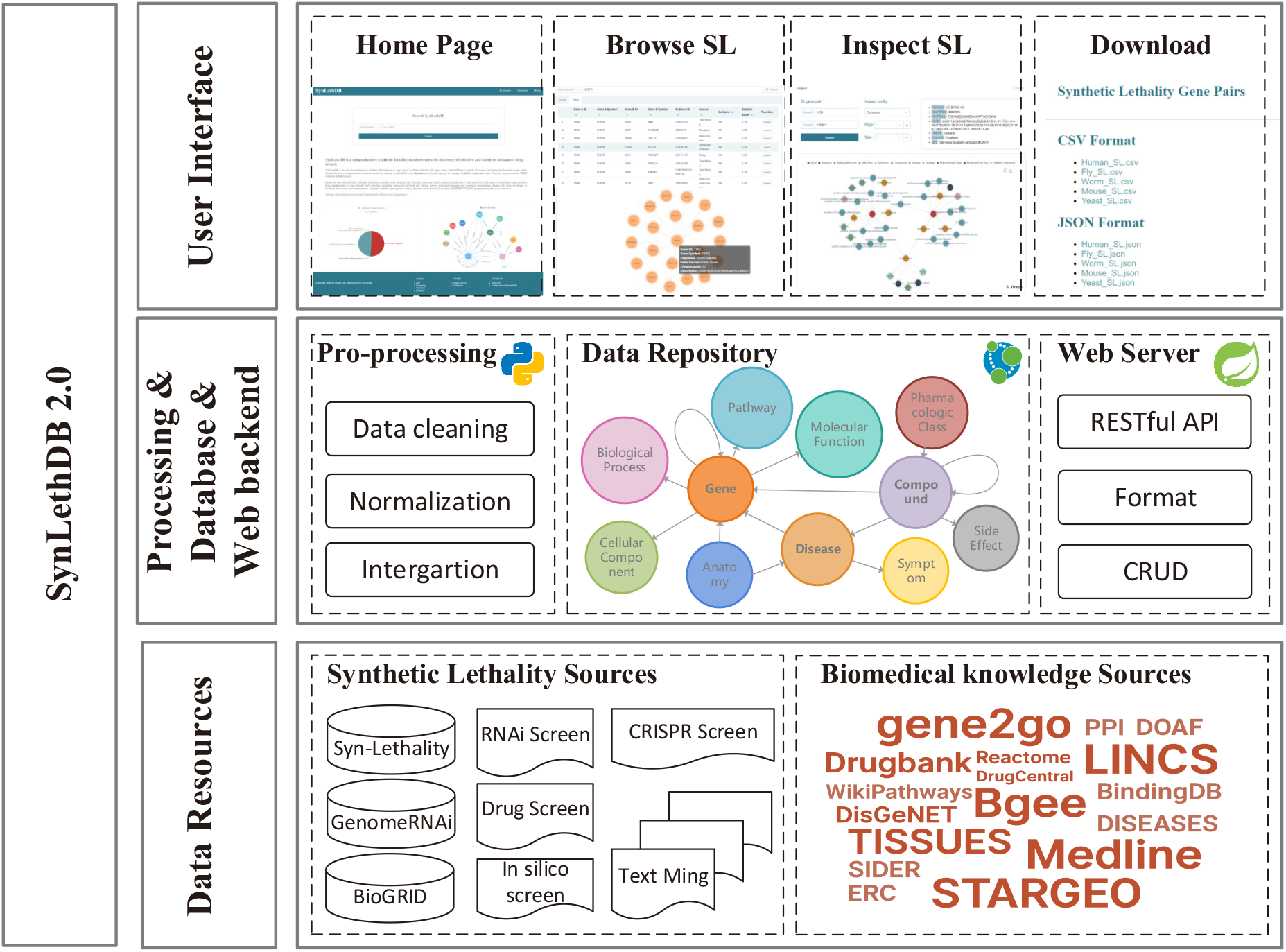
Architecture of SynLethDB 2.0. The bottom layer shows the data sources of SLs and other biomedical knowledge. The middle layer shows the data pre-processing steps, database storage, and web server. The top layer shows the main functional modules of the user interface.

## Materials and Methods

### Data sources

The new version of SynLethDB contains 50,868 SL pairs which include 35,943 of *Homo sapiens*, 381 of *Mus musculus*, 439 of *Drosophila melanogaster*, 105 of *Caenorhabditis elegans*, and 14,000 of *Saccharomyces cerevisiae*. The first source of the new SL pairs is the research papers on identifying SLs via wet-lab experiments. Using the “synthetic lethal” as a keyword for searching in PubMed, 293 related papers published during years from 2015 to 2019 were extracted for further manual collection of new SL pairs. The second source is public databases containing SL data such as GenomeRNAi [38] and BioGRID [32]. The third source is the SL pairs predicted from wet-lab screen data by computational methods such as GEMINI [50]. For each SL pair, we annotated its species, references to PubMed as supporting evidence, data source type, cell lines and confidence score. Synthetic rescue means mutation in one gene rescues the cell from lethality or growth defect caused by a mutation in another gene [18]. It is related to drug resistance [12] and can be seen as the opposite relationship to SL. We collected 16,207 synthetic rescue (SR) gene pairs and 5,798 non-synthetic lethal (non-SL) gene pairs from the above three sources, which can be used as negative samples to train SL prediction models. Non-synthetic lethal pairs could be SR or other relationships. Some gene pairs show up in both SL and non-SL datasets, depending on the different cell lines or cancer types.

In addition to the above these three kinds of gene pairs, we added 24 types of relationships between genes and other entities (e.g., drugs and cancers). These relationships include gene-compound associations, gene-cancer associations, and other features about genes, cancers and drugs. We manually obtained a list of 53 cancers and curated these relationships from public databases with Python scripts from the open-source project of Hetionet [17]. First, we used the Python script from Hetionet to collect the relationships from data sources. Hetionet collects the relationships between genes, drugs and diseases. We added the relationships among GO terms, pathways and SL genes into the dataset. Every type of relationship was processed into an independent CSV file at first, and then integrated into the Neo4j database for persistent storage with the package Py2Neo. Finally, we constructed a knowledge graph to describe human SL gene pairs and the other 26 types of relationships, named SynLethKG.

### Data quality improvement

In addition to collecting the data about SLs, we have also improved the annotation quality of SL gene pairs. First, we collected the SL entries from different sources into one TSV format file to facilitate subsequent unified processing. Second, we completed the missing identifiers of the genes. With annotation packages from Bioconductor, which provide genome annotations for different species, we completed the missing Entrez ID of a gene by its gene symbol or completed the missing gene symbol by its Entrez ID. Third, we deleted entries that still lacked gene IDs or gene symbols. These entries lacked gene IDs or gene symbols because they contained incorrect gene symbols or IDs, which may be due to recording errors from the original sources. After that, we downloaded the latest version of the gene annotations from the Gene Entrez database [30] on the NCBI FTP site, then deleted the SL entries that contain genes deprecated by the current NCBI Gene Entrez database. Lastly, we removed duplicate SL entries that have the same genes and PubMed IDs. The SL entries that contain the gene SL pair but were from different sources, are merged into one entry.

Furthermore, unlike in the first version of the database where SLs were stored in the form of records in a table, in the new graph database SLs are stored as undirected edges between two gene nodes. Hence, only one SL entry can be stored between a pair of genes. The species, references to PubMed, supporting evidence, cell lines, and other relevant information about an SL entry are stored as properties of the edge, and the gene annotation information is stored as the node properties.

### Construction of graph database

In the previous version of SynLethDB, we used the relational database management system, MySQL, to store the data. In this version, we chose to use a graph database system, Neo4j, to store SL pairs and related biomedical knowledge. Graph database is more suitable for many-to-many relationships. The relational database computes the relationships at query time through expensive operations such as JOIN. By contrast, the graph database stores the relationships as edges which processes and queries the relationships more efficiently. We used the Java framework of Spring Data Neo4j, as middle-ware for object-graph mapping and data persistence. All the queries are accessible to users through the front-end interface in the form of Representational State Transfer (REST) API using Hypertext Application Language (HAL) as the media type.

The front end of SynLethDB is a single-page application built using VueJS and Element UI. When changing the tabs, only the required content is updated instead of the whole page, enabling faster responses. It allows us to cache searching queries from users and create a better user experience until the web session is updated. Interactive and expandable graph viewers are developed with the ECharts JavaScript library to visualize the query results as connections in the graph database.

We used Nginx as a reverse proxy to hide the real host of SynLethDB for web security. In the deployment of SynLethDB, we followed the micro-services architecture to get a higher scalability and reduce downtime through fault isolation. The database, web-interface, and web server are all hosted in independent docker containers and arranged by Docker Compose. These services can be easily migrated, automatically deployed, and quickly restored, which ensures high accessibility of SynLethDB.

### Confidence scores of SL pairs

The SLs in our database were collected from different sources, including manually checked publications, existing databases, computational predictions and text mining. According to the types of sources, we use a two-step strategy, i.e., quantification and integration, to calculate the final confidence scores, following the strategy of SynLethDB 1.0 [13]. The main differences from the previous version are the individual scores in the quantification and weight factors in the integration.

In the quantification step, the quantitative score is assigned based on the experimental methods that were used, and a individual score is assigned for each kind of evidence. To incorporate the new source of CRISPR screening, we reset the individual scores as Table 1 shows.

**Table 1.** Quantitative scores assigned to SLs according to experimental methods.

| Method | Score |
| --- | --- |
| CRISPR interference | 0.85 |
| Drug inhibition | 0.75 |
| RNA interference | 0.75 |
| Low-throughput | 0.80 |
| High-throughput | 0.50 |

In the integration step, we integrated the scores of different types of sources into a normalized confidence score for every SL pair. Different weights are assigned according to the source types. The default weights for biochemical experiment, existing databases, computational prediction and text mining are 0.8, 0.5, 0.3 and 0.2, respectively. The weights are set empirically, and users can customize these values according to their own experience or needs when they browse the SLs on the web interface of our database.

### Gene set enrichment analysis

Given a gene *g*, let *G* denote the set of all SL partner genes of *g*. The enrichment analysis is to find out the pathways and GO terms from each of the three ontologies (i.e., biological process, molecular function and cellular component) that occur significantly more frequently than random in the gene set *G*. We implemented two enrichment analysis methods based on the degree information and p-value respectively.

#### Degree-based gene set enrichment analysis

An SLPR score inspired by PageRank [33] was computed for each pathway or GO term associated with the gene set *G*. The pathways and GO terms can be ranked based on their SLPR scores. A larger SLPR score means that a pathway or GO term is more closely associated with the gene set. The SLPR score is defined as:

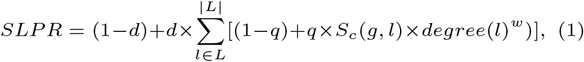

where *d* is a damping factor set to 0.85, *q* is another damping factor set to 0.8, and *w* is set to *−*1 to reflect a negative correlation. For a specific pathway or GO term, *L* represents the subset of genes in set *G* that are directly connected with it. Given a gene *l ∈ L, S_c_*(*g, l*) is the confidence score of the SL pair (*g, l*), and *degree*(*l*) is the number of pathways or GO terms associated with *l*.

#### P-value-based gene set enrichment analysis

Assume that *M* is the number of genes in *G* and *N* is the number of genes having SL partners in the whole database. Given a specific pathway or GO term, *n* is the total number of genes associated with it and *m* is the number of genes in *G* associated with it. To show the enrichment of the gene set *G* with the pathway or GO term, we calculate a p-value as follows [21].

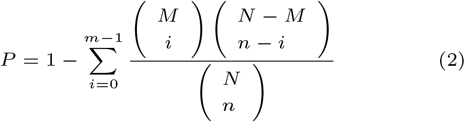

Thus, we attain a list of pathways or GO terms sorted in order of the p-values. A smaller p-value means that *G* is more enriched with the given pathway or GO term.

### SynLethDB 2.0 portal

A user-friendly web interface has been developed for SynLethDB to facilitate data visualization, analysis and interpretation. Compared with the first version, SynLethDB 2.0 is more user friendly in that it provides more interactive searching options and network-view of relationships with state-of-art web design. On the website of SynLethDB, we provide a general introduction to the database, as well as the search bar for looking up SLs by gene symbols or gene IDs. Other functionalities of SynLethDB can be accessed by menu tabs on the website as follows.

#### Searching and browsing the SLs

In the first version of SynLethDB, users could only search for SLs by genes. In this new version, we collected 14,116 gene-cancer relationships and 56,921 gene-compound relationships for those genes involved in SLs from DisGeNET [34], Drugbank [47], and BindingDB [10]. Based on these new data, we offer two new options for searching, namely, “search SL by disease” and “search SL by compound”, and provide the auto-complete function to the list of all available cancers or compounds in SynLethDB. The searching results are shown in a table viewer.

#### Customizable confidence scores for SLs

A confidence score reflects an SL’s credibility based on its sources, which can be used to rank SLs. As mentioned earlier, we use a two-step scoring procedure (i.e., quantification and integration) to assign a confidence score based on the sources of the SL. In the quantification step, we assigned the quantitative scores to SL pairs according to their experimental methods as shown in Table 1. In the integration step, we provide default values for the weight factors, but allow users to customize these weights to facilitate them to extract the SLs of a certain type of source that they are most interested in. When searching and browsing SLs by genes, users can adjust the weight factors of source types and rank results by the confidence scores.

#### Searching and browsing the knowledge graph SynLethKG. SynLethKG

contains relationships that describe various features for genes, cancers and drugs. With the “Inspect SL” functionality, all these relationships are categorized by their node types and can be browsed through an interactive graph viewer. Starting with SL genes to be inspected, users only need to click on the node they are about to inspect, and the graph viewer can fetch and visualize the results. The type of relationships and the number of edges to be displayed can be specified by the users. Properties of the nodes and edges, such as data sources and entity IDs, can also be viewed through an infobox in the upper right corner.

#### Gene set enrichment analysis of SL partners

We developed two methods for gene set enrichment analysis based on p-values and node degrees respectively. Both methods take a gene symbol as input, and conduct gene set enrichment analysis for the SL partners of this gene. The output includes the rank of the pathways and GO terms separately. A higher ranking of a pathway or a GO term indicates that the SL partners of this gene is more enriched with this pathway or GO term. The p-value-based enrichment analysis tool ranks the results by p-value calculated in Equation 2, and a lower p-value corresponds to higher ranking. Meanwhile, the degree-based enrichment analysis tool ranks the pathways and GO terms based on the SLPR score as calculated in Equation 1, and a higher SLPR score corresponds to a higher ranking.

#### Data access and download

We provide a download page to make it easy for users to retrieve a large amount of data. All the SL gene pairs are classified by species and can be downloaded in either CSV or JSON format. We provide the files of SynLethKG in the formats of CSV, JSON, and GraphML for users to download. In particular, the datasets in GraphML format can be imported to other software tools such as Gephi and Cytoscape for analysis and visualization. For users who prefer the triplet format, we also provide a CSV file that contains all the relationships in the format (source, relationship, target). All the data can be freely accessed and downloaded without a login requirement. RESTful APIs are also provided for users to access and analyze the data by running the scripts in programming languages such as Python and R.

#### User manual

To lower the learning curve for new users of SynLethDB, we offer a web page containing a user manual, which gives an introduction to every functionality of SynLethDB, as well as examples of using the web interface and the RESTful APIs.

## Results

### Comparison with other databases

In this subsection, we compare SynLethDB 2.0 with SynLethDB 1.0 [13] and Synthetic Lethality Knowledge Graph (SLKG) [51]. From the comparison, we observe that SynLethDB 2.0 excels in the following aspects.

First, SynLethDB 2.0 is the most up-to-date and most comprehensive database for SLs. SynLethDB 2.0 contains 50,868 SL pairs in total, almost doubling the number of SL pairs in SynLethDB 1.0. In particular, SynLethDB 2.0 contains 35,943 human SLs, 381 mouse SLs, 439 fly SLs, 14,000 yeast SLs and 105 worm SLs as shown in Table 2. Regarding human SLs, SynLethDB 1.0 and SLKG have comparable numbers of SL pairs, while the number of SynLethDB 2.0 is almost 1.8 times that of each of them. Similar to SynLethDB 1.0, we also provide the HGNC gene symbols, Entrez gene IDs, PubMed IDs of its original publications, types of sources and the confidence score calculated according to the sources for each SL pair in SynLethDB 2.0. Note that we updated the confidence scores by considering new sources of SLs such as CRISPR screening and allowing user-defined weight factors.

**Table 2.**
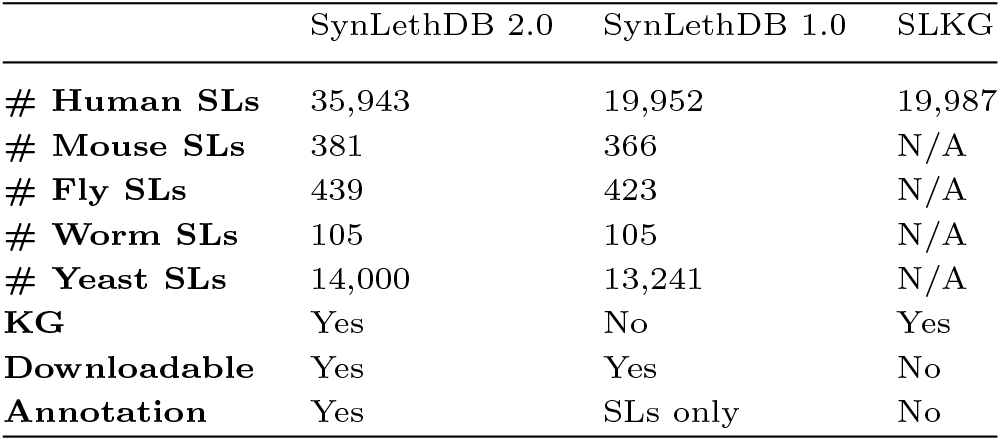
Comparison of statistics among existing databases.

|  | SynLethDB 2.0 | SynLethDB 1.0 | SLKG |
| --- | --- | --- | --- |
| # Human SLs | 35,943 | 19,952 | 19,987 |
| # Mouse SLs | 381 | 366 | N/A |
| # Fly SLs | 439 | 423 | N/A |
| # Worm SLs | 105 | 105 | N/A |
| # Yeast SLs | 14,000 | 13,241 | N/A |
| KG | Yes | No | Yes |
| Downloadable | Yes | Yes | No |
| Annotation | Yes | SLs only | No |

Second, SynLethDB 2.0 provides more types of biomedical knowledge. SLKG [51] is a knowledge graph which contains the relationships among genes, drugs, and cancers. SynLethKG in SynLethDB 2.0 contains much more types of entities and relationships, including biological processes, pathways, molecule functions and cellular components for genes, pharmacologic classes and side effects for drugs, symptoms and anatomies for cancers. Overall, there are 37,341 entities/nodes and 1,405,652 relationships/edges in SynLethKG as shown in Table 3 clearly, SynLethDB 2.0 provides a more comprehensive knowledge graph about human SLs than SLKG. The types of relationships and their numbers are listed in Table 4. In addition, SynLethDB 2.0 retains and corrects the annotations of SLs in SynLethDB 1.0, and also adds annotations to nodes and edges in SynLethKG, such as the name of entity, the data source, and the link to entity in the original data source, and other annotations such as the organisms of genes and the thresholds used when extracting the relationships. Therefore, SynLethDB 2.0 provides more comprehensive annotations for the entries and relationships. Table 5 shows the number of each type of entities in SynLethKG, and the average numbers of annotations and relationships of each kind of entities. The average number of relationships for each type of nodes is counted by adding the number of edges among all the nodes and dividing the sum by the number of nodes.

**Table 3.**
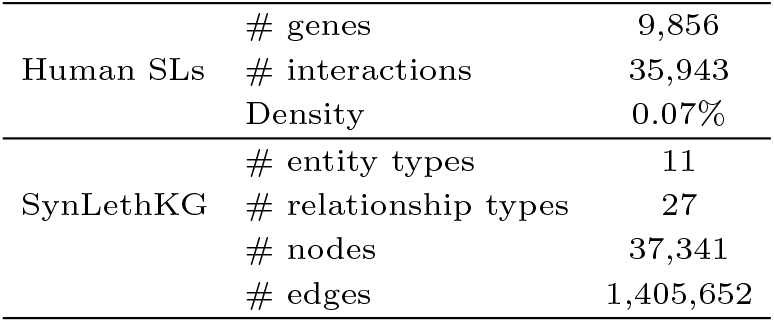
Statistics about the knowledge graph SynLethKG

**Table 4.** NUmbers of the relationships in SynLethKG.

| Type | # Edges |
| --- | --- |
| (Anatomy, downregulates, Gene) | 31 |
| (Anatomy, expresses, Gene) | 358,005 |
| (Anatomy, upregulates, Gene) | 26 |
| (Compound, binds, Gene) | 11,453 |
| (Compound, causes, Side Effect) | 135,063 |
| (Compound, downregulates, Gene) | 17,506 |
| (Compound, palliates, Cancer) | 42 |
| (Compound, resembles, Compound) | 5,500 |
| (Compound, treats, Cancer) | 282 |
| (Compound, upregulates, Gene) | 13,573 |
| (Cancer, associates, Gene) | 7,708 |
| (Cancer, downregulates, Gene) | 988 |
| (Cancer, localizes, Anatomy) | 1,444 |
| (Cancer, presents, Symptom) | 1,048 |
| (Cancer, resembles, Cancer) | 106 |
| (Cancer, upregulates, Gene) | 1,263 |
| (Gene, covaries, Gene) | 16,985 |
| (Gene, interacts, Gene) | 87,103 |
| (Gene, non-synthetic lethal, Gene) | 2,831 |
| (Gene, participates, Biological Process) | 393,049 |
| (Gene, participates, Cellular Component) | 59,054 |
| (Gene, participates, Molecular Function) | 65,207 |
| (Gene, participates, Pathway) | 41,790 |
| (Gene, regulates, Gene) | 147,639 |
| (Gene, synthetic lethal, Gene) | 35,943 |
| (Gene, synthetic rescue, Gene) | 895 |
| (Pharmacologic Class, includes, Compound) | 1,118 |

**Table 5.** Statistics about the entities in SynLethKG.

| labels(n) | Size | Avg_Ann<sup>1</sup> | Avg_Rel<sup>2</sup> |
| --- | --- | --- | --- |
| SideEffect | 5,664 | 5.00 | 23.85 |
| Gene | 14,100 | 8.00 | 112.99 |
| BiologicalProcess | 12,141 | 5.00 | 32.37 |
| Compound | 1,898 | 7.00 | 100.12 |
| MolecularFunction | 3,012 | 5.00 | 21.65 |
| Anatomy | 390 | 6.64 | 921.81 |
| CellularComponent | 1,619 | 5.00 | 36.48 |
| Pathway | 2,069 | 5.00 | 20.63 |
| Symptom | 325 | 5.00 | 3.224 |
| PharmacologicClass | 357 | 6.00 | 3.13 |
| Cancer | 53 | 5.00 | 245.04 |

<sup>1</sup>The average number of annotations of each type of nodes.
<sup>2</sup>The average number of relationships of each type of nodes.

Third, the data in SynLethDB 2.0 are freely downloadable. For users who intend to do in-depth data analysis, a local copy of data would be necessary. SynLethDB 2.0 allows users to download the datasets from the website, while SLKG only allows the query results to be downloaded.

### SynLethKG for cancer driver genes

In SynLethKG, we collected various relationships for the genes involved in the SL pairs, including gene-gene relationships (gene expression covariation, gene interaction and gene regulation), Gene Ontology (GO) annotations and pathways as shown in Table 4. In particular, SynLethKG has 14,100 genes, 12,141 biological processes, 3,012 molecular functions, 1,619 cellular components and 2,026 pathways etc. as nodes and their relationships as edges in Table 5.

Moreover, SynLethKG also contains 9,959 relationships between the genes and 53 cancers from DisGeNET database [34] and 42,532 relationships between the genes and 1,898 compounds from DrugBank database [47]. For cancers, 325 symptoms and 390 anatomies are included as entities to describe the cancer features. For drugs, 357 pharmacologic classes and 5,664 side-effects are included as entities to describe drug features. Based on the same strategy as in SL-BioDP [8], we counted the numbers of cancer driver genes and genes from hallmark cancer pathways in 32 cancer types from TCGA contained in SynLethKG, as well as the numbers of their SL partners and related drugs.

Figure 2 shows the numbers of cancer drive genes, their SL partners and related drugs. We can observe that several cancers have quite a few SL pairs and drugs related to their driver genes, including BLCA, BRCA, CESC, COADREAD, HNSC, LGG, LIHC, SKCM and UCEC. Figure 2 demonstrates that our database contains useful information about many genes, SLs and drugs related to cancers, which could be potentially used as an interpretation tool for data-driven SL-based drug target discovery.

**Fig. 2.**
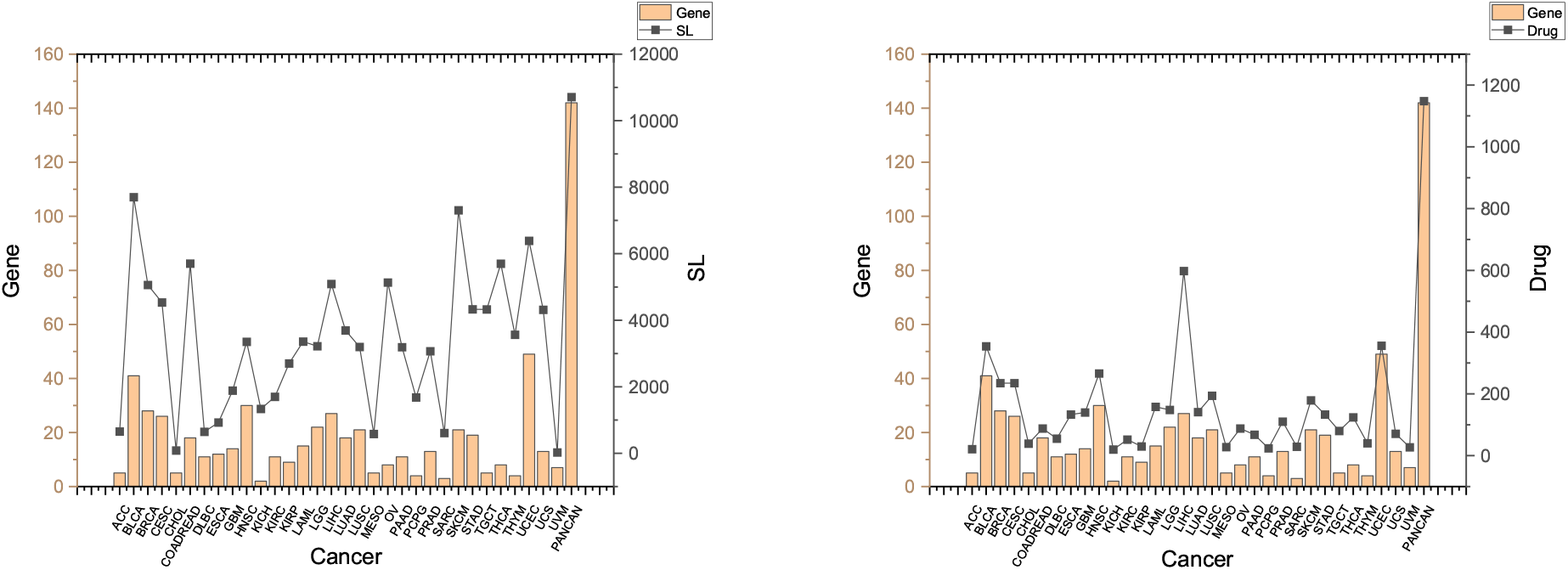
SLs and drugs in SynLethKG for driver genes of 32 cancers and pan-caner. The bar chart shows the numbers of cancer driver genes in SynLethKG. In the left figure, the line chart represents the numbers of SLs containing the cancer driver genes in SynLethKG, and in the right figure, the line chart represents the numbers of drugs associated with the driver genes in SynLethKG.

### SynLethKG for predictive modeling

Based on known SL data and related knowledge, new SLs can be predicted. Compared to the previous version, SynLethDB 2.0 provides more SL data and related knowledge as features to allow more accurate prediction of SLs. In the work of Wang et al. [45], it was demonstrated that additional information from the SynLethKG could guide a model to achieve better performance. By extracting gene interaction, gene regulation and gene co-variation relationships from SynLethKG we constructed a sub-graph of the SynLethKG knowledge graph, named G2G graph. We compare the SL prediction performances of models trained against the SL dataset, G2G graph and SynLethKG to analyze the contributions of different types of knowledge to SLs prediction.

We used all 35,943 human SLs as positive samples and randomly up-sampled the negative samples from 3,726 synthetic rescue (SR) and non-synthetic lethal (non-SL) pairs to the same size of the positive samples to avoid the influence of data imbalance. The dataset is randomly split into a training set and test set, constituting 70% and 30% of the original dataset, respectively. The training set is used to train models, and the associated test set is used to evaluate the trained model. In particular, we built the traditional machine learning model of Random Forest (RF), and a graph neural network, Graph Convolutional Network (GCN), for SL prediction.

The classical Random Forest model, with 30 estimators and maximum depth 10, was implemented using Scikit-Learn, a Python library for machine learning. We extracted 9 features from the gene graphs for every SL pair, which were calculated by a graph algorithm library in Neo4j named “Graph Science Data” from three aspects, i.e. common neighbors, triangles and clustering coefficients, and community detection. The features based on common neighbors include the number of total neighbors, the number of common neighbors and the preferential attachment value. The features based on triangles and clustering coefficients include the maximum and minimum numbers of triangles that a node is part of, and the maximum and minimum values of clustering coefficients which show if the neighbors are also connected. Two features extracted from community detection are the Boolean values (i.e. represented by 0 or 1) about whether a pair of nodes are in the same community detected by label propagation algorithm [35] and Louvaion algorithm [28]. We applied RF on three graphs, namely, G2G graph, SL graph and SL+G2G graph, to test the contribution of knowledge graph to SL prediction. SL+G2G graph comprises nodes and edges from either the SL graph or the G2G graph. We did not run the RF model on SynLethKG because the RF model requires pre-processing the nine kinds of features for each type of the relationships in SynLethKG before training. Thus, the performance on SynLethKG is demonstrated by the GCN model.

The graph neural network model, GCN, was implemented by a Python library named Deep Graph Library (DGL) [43], with two graph convolution layers. To further test the performance of the model on a dataset with more knowledge, we tested the performance of the GCN model on the G2G graph and the SynLethKG graph, with the same pre-processing of training set and testing set. There are 24 kinds of relationships in the features of SynLethKG graph instead of 27, because the SLs used as positive samples and the SRs and non-SLs used as negative samples and should not be counted as features when training the model to avoid leaking label information in training data.

The performances of the models are measured by F1, AUROC and AUPR scores. As Table 6 shows, the RF model trained against the SL graph can perform better than the RF model trained anainst the G2G graph. However, the combination of SL and G2G can achieve even better performance. This indicates that the features extracted from other relationships can contribute to SL prediction. The GCN also has better performance on the SynLethKG dataset (as shown in the last row of Table 6) than on the G2G dataset, which indicates that adding more knowledge from SynLethKG can further improve the performance of SL prediction.

**Table 6.** Performance of SL prediction by two models.

| Model | Dataset | AUROC | AUPR | F1 |
| --- | --- | --- | --- | --- |
| RF | G2G | 0.873±0.003 | 0.912±0.002 | 0.867±0.003 |
| RF | SL | 0.938±0.002 | 0.959±0.002 | 0.927±0.002 |
| RF | SL+G2G | 0.938±0.002 | 0.965±0.001 | 0.934±0.002 |
| GCN | G2G | 0.862±0.003 | 0.891±0.002 | 0.875±0.003 |
| GCN | SynLethKG | 0.870±0.029 | 0.898±0.016 | 0.886±0.021 |

### Case study

To demonstrate how to use SynLethDB to discover drug targets, let us do a case study of searching SL partners of BRCA1 in breast cancer through the web interface as shown in Figure 3. First, with the “Search SL by disease” module in SynLethDB, we choose “breast cancer” as the disease and select the relationship “Disease Associates Gene”. Then, the first line of the results show that breast cancer is associated with BRCA1. By clicking the “Search SL” button in the function column, we searched the SL partners of BRCA1. The result shows that PARP2 is an SL partner of BRCA1 with a high confidence score (0.87). Here we choose this SL pair for further inspection. After clicking the “Inspect” button in the function column, we can browse more knowledge about this SL pair. Different types of biomedical relationships can be browsed by clicking the nodes in the graph. For example, we can see that both BRCA1 and PARP2 are associated with the “ovarian cancer” and “breast cancer” diseases, and BRCA1 participates in the “DNA Damage Response” pathway. As breast cancer down-regulates BRCA1 and BRCA1 has an SL partner, PARP2, we check out compounds that down-regulate PARP2 as candidate drugs for breast cancer. As shown in the graph, Rucaparib, Talazoparib, Niraparib, and Olaparib all bind with PARP2. On the other hand, we can also search Rucaparib, Talazoparib, Niraparib, and Olaparib from the “Search SL by compound” page. In this way, we can find that PARP2 is a drug target. Through this case study, we show the basic functionalities of the database through the web interface, which can be used to explore potential anti-cancer drug targets based on SL or analyze the biological mechanisms behind SLs.

**Fig. 3.**
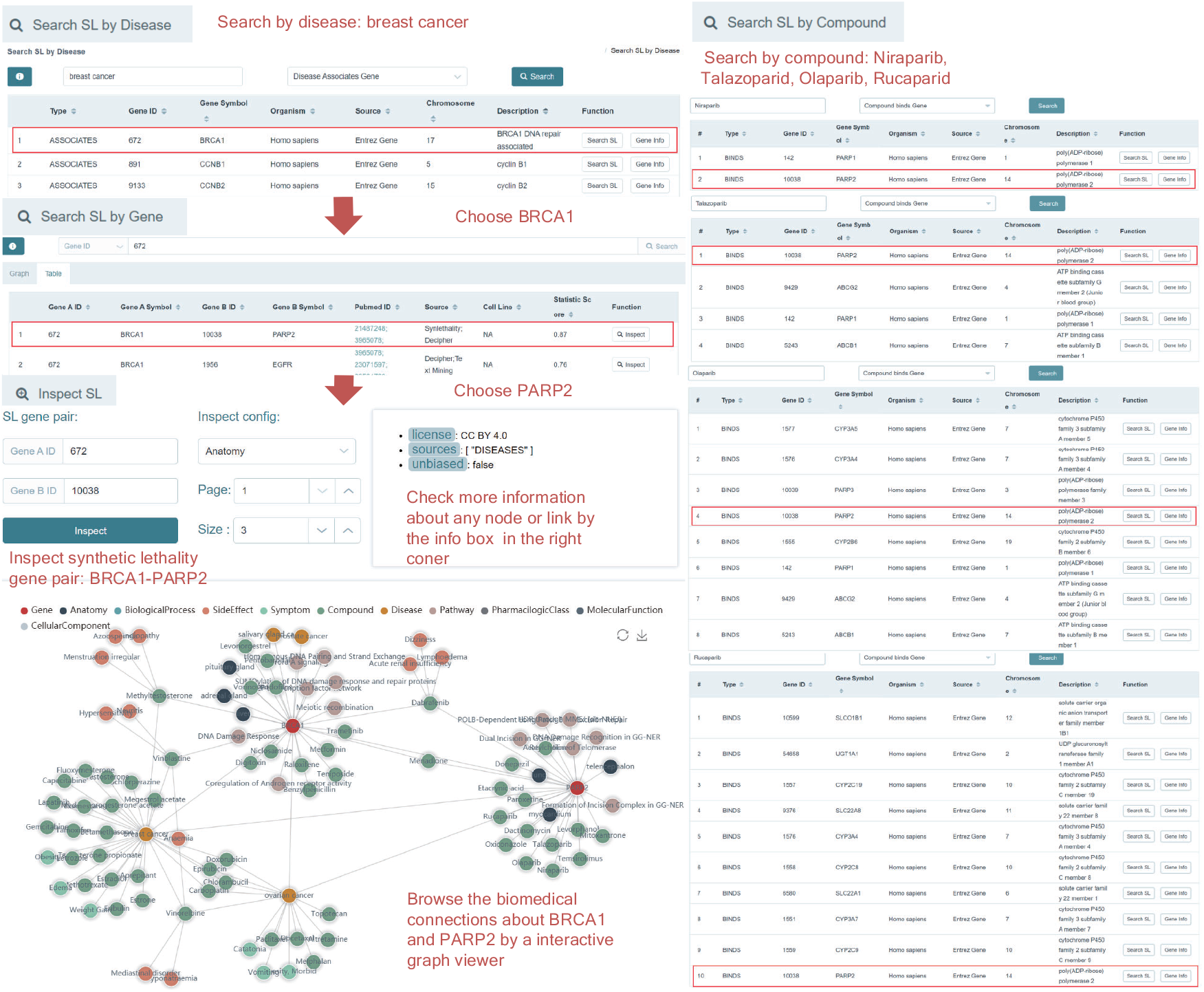
The case study on BRCA1 in breast cancer. Using “Search SL by disease”, the SL genes associated with the disease will be shown. BRCA1 is a gene that has SL partners and is down-regulated in breast cancer. Click the “Search SL” button, and it shows that PARP2 is an SL partner of BRCA1 with a high confidence score (0.87). Inspecting this pair of SL genes, we notice that BRCA1 and PARP2 are both associated with the “ovarian cancer” and “breast cancer” diseases, and BRCA1 participates in the “DNA Damage Response” pathway. The result that breast cancer down-regulates BRCA1 and PARP2 is an SL partner of BRCA1 indicates that PARP2 is a drug target for breast cancer. Rucaparib, Talazoparib, Niraparib, and Olaparib all bind with PARP2. Using “Search SL by compound”, we also identify PARP2 as a target gene of Rucaparib, Talazoparib, Niraparib and Olaparib.

## Discussion and Conclusion

With the development of RNAi and CRISPR screening technologies, data about synthetic lethality have increased rapidly in the past few years. We have been continuously collecting SL data, integrating them into SynLethDB and improving the annotation quality. In this version, we have integrated more biomedical knowledge about human SLs into a knowledge graph called SynLethKG. The additional knowledge can provide more features for SL prediction and improve the performance of the predictive model. A similar procedure can be applied to predicting drugs based on SLs. We also provided a new web interface with online services for data browsing, visualization and analysis. For instance, “Search SL by disease” can facilitate SL-based cancer drug discovery. The “SL inspect” functionality displays relationships between a pair of SL genes from multiple sources in one intuitive graph. Enrichment analysis tools help analyze the most relevant pathways and GO of a gene’s SL partners. SynLethDB has been used as a source of training or testing datasets by many computational methods for SL prediction, and this new version of SynLethDB provides a larger and more comprehensive dataset for these methods. In addition, we realize that the data in SynLethDB are enriched with SLs of some hub genes, such as KRAS, because they are more experimentally studied. This kind of data skewness may introduce some bias, which makes a model learn superficial patterns and achieve inflated performance.

The overall goal of SynLethDB is to increase the understanding of SL mechanisms and to facilitate drug discovery. In the future, we will continue to collect new SLs and enhance the functionalities of the database. For instance, we will add genomics data and cell line annotations to make SLs more context-specific. In addition, we can create more efficient path queries based on the graph database to find the pathways shared between SL pairs and interactions between SLs and drugs.

## Key Points

- To provide the scientific community with the latest data about SL gene pairs, we developed SynLethDB 2.0. The number of SLs included has increased from 34,089 to 50,868, including new results from CRISPR screening.
- We have integrated 27 relationships of 11 entities and constructed a knowledge graph about human SL gene pairs, named SynLethKG, to provide more biomedical knowledge about SLs.
- SynLethKG contains comprehensive knowledge about SLs and provides useful features for SL prediction. Similarly, the relationships between SLs, drugs and cancers can also be used for drug discovery.
- SynLethDB 2.0 provides a new website interface to facilitate the retrieval and visualization of knowledge graph. Online tools have also been developed, including drug-based and cancer-based SL searching, p-value-based and degree-based gene enrichment analysis of SL partners, and the display of the subgraph between a pair of SL genes.
- The data and knowledge graph SynLethKG contained in SynLethDB 2.0 can be fully accessed and downloaded through the website or RESTful APIs.

## Data Availability

The datasets of SynLethDB 2.0 are freely available at http://synlethdb.sist.shanghaitech.edu.cn/v2.

## Author contributions statement

JZ, MW and HL conceived the study. JW and SZ collected the data and performed the analysis. JW, XH, and LW developed the SynLethKG knowledge graph and the SynLethDB database. JW drafted the manuscript with critical input from JZ and MW. All authors reviewed the manuscript.

**Jie Wang** is a master student at School of Information Science and Technology, ShanghaiTech University, Shanghai, China.

**Min Wu** is a research scientist at the Data Analytics Department, Institute for Infocomm Research (I2R) under the Agency for Science, Technology and Research (A*STAR), Singapore. His current research interests include machine learning, data mining, and bioinformatics.

**Xuhui Huang** is a master student at National University of Singapore with Specialisation in Artificial Intelligence.

**Li Wang** is a research assistant at the School of Information Science and Technology, ShanghaiTech University, Shanghai, China. Her research interests include bioinformatics and medical image processing.

**Sophia Zhang** is a graduate of School of Agriculture and Life Science, Cornell University, USA.

**Hui Liu** is a professor at School of Computer Science and Technology, Nanjing Tech University, Nanjing, China. His research interests include the anti-cancer drug screening by means of bioinformatics and deep learning.

**Jie Zheng** is an associate professor at the School of Information Science and Technology, ShanghaiTech University, Shanghai, China. His research interests include bioinformatics, biomedical data science, artificial intelligence, precision medicine and AI for drug discovery.

## References

1. Steven R Bartz, Zhan Zhang, Julja Burchard, Maki Imakura, Melissa Martin, Anthony Palmieri, Rachel Needham, Jie Guo, Marcia Gordon, Namjin Chung, et al. Small interfering RNA screens reveal enhanced cisplatin cytotoxicity in tumor cells having both BRCA network and TP53 disruptions. Molecular and Cellular Biology, 26(24):9377–9386, 2006.

2. Jonathan L Blank, Xiaozhen J Liu, Katherine Cosmopoulos, David C Bouck, Khristofer Garcia, Hugues Bernard, Olga Tayber, Greg Hather, Ray Liu, Usha Narayanan, et al. Novel DNA damage checkpoints mediating cell death induced by the NEDD8-activating enzyme inhibitor MLN4924. Cancer Research, 73(1):225–234, 2013.

3. Helen E Bryant, Niklas Schultz, Huw D Thomas, Kayan M Parker, Dan Flower, Elena Lopez, Suzanne Kyle, Mark Meuth, Nicola J Curtin, and Thomas Helleday. Specific killing of BRCA2-deficient tumours with inhibitors of poly (ADP-ribose) polymerase. Nature, 434(7035):913–917, 2005.

4. Ruichu Cai, Xuexin Chen, Yuan Fang, Min Wu, and Yuexing Hao. Dual-dropout graph convolutional network for predicting synthetic lethality in human cancers. Bioinformatics, 36(16):4458–4465, 2020.

5. Jan-Gowth Chang, Chia-Cheng Chen, Yi-Ying Wu, Ting-Fang Che, Yi-Syuan Huang, Kun-Tu Yeh, Grace S Shieh, and Pan-Chyr Yang. Uncovering synthetic lethal interactions for therapeutic targets and predictive markers in lung adenocarcinoma. Oncotarget, 7(45):73664, 2016.

6. XiuLiang Cui, Lu Han, Yang Liu, Ying Li, Wen Sun, Bin Song, Wenxia Zhou, Yongxiang Zhang, and Hongyang Wang. siGCD: a web server to explore survival interaction of genes, cells and drugs in human cancers. Briefings in Bioinformatics, 2021.

7. Shaoli Das, Xiang Deng, Kevin Camphausen, and Uma Shankavaram. DiscoverSL: an R package for multi-omic data driven prediction of synthetic lethality in cancers. Bioinformatics, 35(4):701–702, 2019.

8. Xiang Deng, Shaoli Das, Kristin Valdez, Kevin Camphausen, and Uma Shankavaram. Sl-biodp: Multi-cancer interactive tool for prediction of synthetic lethality and response to cancer treatment. Cancers, 11(11):1682, 2019.

9. Th Dobzhansky. Genetics of natural populations. XIII. Recombination and variability in populations of Drosophila pseudoobscura. Genetics, 31(3):269, 1946.

10. Michael K Gilson, Tiqing Liu, Michael Baitaluk, George Nicola, Linda Hwang, and Jenny Chong. BindingDB in 2015: a public database for medicinal chemistry, computational chemistry and systems pharmacology. Nucleic Acids Research, 44(D1):D1045–D1053, 2016.

11. Mark A Gregory, Tzu L Phang, Paolo Neviani, Francesca Alvarez-Calderon, Christopher A Eide, Thomas O’Hare, Vadym Zaberezhnyy, Richard T Williams, Brian J Druker, Danilo Perrotti, et al. Wnt/Ca2+/NFAT signaling maintains survival of Ph+ leukemia cells upon inhibition of Bcr-Abl. Cancer Cell, 18(1):74–87, 2010.

12. Yunyan Gu, Ruiping Wang, Yue Han, Wenbin Zhou, Zhangxiang Zhao, Tingting Chen, Yuanyuan Zhang, Fuduan Peng, Haihai Liang, Lishuang Qi, et al. A landscape of synthetic viable interactions in cancer. Briefings in Bioinformatics, 19(4):644–655, 2018.

13. Jing Guo, Hui Liu, and Jie Zheng. SynLethDB: synthetic lethality database toward discovery of selective and sensitive anticancer drug targets. Nucleic Acids Research, 44(D1):D1011–D1017, 2016.

14. Kyuho Han, Edwin E Jeng, Gaelen T Hess, David W Morgens, Amy Li, and Michael C Bassik. Synergistic drug combinations for cancer identified in a CRISPR screen for pairwise genetic interactions. Nature Biotechnology, 35(5):463–474, 2017.

15. Leland H Hartwell, Philippe Szankasi, Christopher J Roberts, Andrew W Murray, and Stephen H Friend. Integrating genetic approaches into the discovery of anticancer drugs. Science, 278(5340):1064–1068, 1997.

16. Andreas Heinzel, Maximilian Marhold, Paul Mayer, Michael Schwarz, Erwin Tomasich, Arno Lukas, Michael Krainer, and Paul Perco. Synthetic lethality guiding selection of drug combinations in ovarian cancer. PLoS ONE, 14(1):e0210859, 2019.

17. Daniel Scott Himmelstein, Antoine Lizee, Christine Hessler, Leo Brueggeman, Sabrina L Chen, Dexter Hadley, Ari Green, Pouya Khankhanian, and Sergio E Baranzini. Systematic integration of biomedical knowledge prioritizes drugs for repurposing. Elife, 6:e26726, 2017.

18. Thomas höfken and Elmar Schiebel. Novel regulation of mitotic exit by the Cdc42 effectors Gic1 and Gic2. The Journal of Cell Biology, 164(2):219–231, 2004.

19. Yuxuan Hu, Chia-hui Chen, Yang-yang Ding, Xiao Wen, Bingbo Wang, Lin Gao, and Kai Tan. Optimal control nodes in disease-perturbed networks as targets for combination therapy. Nature Communications, 10(1):1–14, 2019.

20. Jiang Huang, Min Wu, Fan Lu, L. Ou-Yang, and Zexuan Zhu. Predicting synthetic lethal interactions in human cancers using graph regularized self-representative matrix factorization. BMC Bioinformatics, 20(19):1–8, 2019.

21. HM James Hung, Robert T O’Neill, Peter Bauer, and Karl Kohne. The behavior of the p-value when the alternative hypothesis is true. Biometrics, pages 11–22, 1997.

22. Livnat Jerby-Arnon, Nadja Pfetzer, Yedael Y Waldman, Lynn McGarry, Daniel James, Emma Shanks, Brinton Seashore-Ludlow, Adam Weinstock, Tamar Geiger, Paul A Clemons, et al. Predicting cancer-specific vulnerability via data-driven detection of synthetic lethality. Cell, 158(5):1199–1209, 2014.

23. William G Kaelin. The concept of synthetic lethality in the context of anticancer therapy. Nature Reviews Cancer, 5(9):689–698, 2005.

24. William G Kaelin et al. Choosing anticancer drug targets in the postgenomic era. The Journal of Clinical Investigation, 104(11):1503–1506, 1999.

25. Joo Sang Lee, Avinash Das, Livnat Jerby-Arnon, Rand Arafeh, Noam Auslander, Matthew Davidson, Lynn McGarry, Daniel James, Arnaud Amzallag, Seung Gu Park, et al. Harnessing synthetic lethality to predict the response to cancer treatment. Nature Communications, 9(1):1–12, 2018.

26. Herty Liany, Anand Jeyasekharan, and Vaibhav Rajan. Predicting synthetic lethal interactions using heterogeneous data sources. Bioinformatics, 36(7):2209–2216, 2020.

27. Yong Liu, Min Wu, Chenghao Liu, Xiao-Li Li, and Jie Zheng. SL2MF: Predicting synthetic lethality in human cancers via logistic matrix factorization. IEEE/ACM Transactions on Computational Biology and Bioinformatics, 17(3):748–757, 2019.

28. Hao Lu, Mahantesh Halappanavar, and Ananth Kalyanaraman. Parallel heuristics for scalable community detection. Parallel Computing, 47:19–37, 2015.

29. Ji Luo, Michael J Emanuele, Danan Li, Chad J Creighton, Michael R Schlabach, Thomas F Westbrook, Kwok-Kin Wong, and Stephen J Elledge. A genome-wide RNAi screen identifies multiple synthetic lethal interactions with the Ras oncogene. Cell, 137(5):835–848, 2009.

30. Donna Maglott, Jim Ostell, Kim D Pruitt, and Tatiana Tatusova. Entrez Gene: gene-centered information at NCBI. Nucleic Acids Research, 39(suppl 1):D52–D57, 2010.

31. Nigel J O’Neil, Melanie L Bailey, and Philip Hieter. Synthetic lethality and cancer. Nature Reviews Genetics, 18(10):613–623, 2017.

32. Rose Oughtred, Chris Stark, Bobby-Joe Breitkreutz, Jennifer Rust, Lorrie Boucher, Christie Chang, Nadine Kolas, Lara O’Donnell, Genie Leung, Rochelle McAdam, et al. The BioGRID interaction database: 2019 update. Nucleic Acids Research, 47(D1):D529–D541, 2019.

33. Lawrence Page, Sergey Brin, Rajeev Motwani, and Terry Winograd. The PageRank citation ranking: Bringing order to the web. Technical report, Stanford InfoLab, 1999.

34. Janet Piñero, Juan Manuel Ramírez-Anguita, Josep Saüch-Pitarch, Francesco Ronzano, Emilio Centeno, Ferran Sanz, and Laura I Furlong. The DisGeNET knowledge platform for disease genomics: 2019 update. Nucleic Acids Research, 48(D1):D845–D855, 2020.

35. Usha Nandini Raghavan, Réka Albert, and Soundar Kumara. Near linear time algorithm to detect community structures in large-scale networks. Physical Review E, 76(3):036106, 2007.

36. Terry Roemer and Charles Boone. Systems-level antimicrobial drug and drug synergy discovery. Nature Chemical Biology, 9(4):222, 2013.

37. Colm J Ryan, Christopher J Lord, and Alan Ashworth. DAISY: picking synthetic lethals from cancer genomes. Cancer Cell, 26(3):306–308, 2014.

38. Esther E Schmidt, Oliver Pelz, Svetlana Buhlmann, Grainne Kerr, Thomas Horn, and Michael Boutros. GenomeRNAi: a database for cell-based and in vivo RNAi phenotypes, 2013 update. Nucleic Acids Research, 41(D1):D1021–D1026, 2013.

39. John Paul Shen, Dongxin Zhao, Roman Sasik, Jens Luebeck, Amanda Birmingham, Ana Bojorquez-Gomez, Katherine Licon, Kristin Klepper, Daniel Pekin, Alex N Beckett, et al. Combinatorial CRISPR–Cas9 screens for de novo mapping of genetic interactions. Nature Methods, 14(6):573–576, 2017.

40. Subarna Sinha, Daniel Thomas, Steven Chan, Yang Gao, Diede Brunen, Damoun Torabi, Andreas Reinisch, David Hernandez, Andy Chan, Erinn B Rankin, et al. Systematic discovery of mutation-specific synthetic lethals by mining pan-cancer human primary tumor data. Nature Communications, 8(1):1–13, 2017.

41. Sriganesh Srihari, Jitin Singla, Limsoon Wong, and Mark A Ragan. Inferring synthetic lethal interactions from mutual exclusivity of genetic events in cancer. Biology Direct, 10(1):1–18, 2015.

42. Zachary Steinhart, Zvezdan Pavlovic, Megha Chandrashekhar, Traver Hart, Xiaowei Wang, Xiaoyu Zhang, Mélanie Robitaille, Kevin R Brown, Sridevi Jaksani, René Overmeer, et al. Genome-wide CRISPR screens reveal a Wnt–FZD5 signaling circuit as a druggable vulnerability of RNF43-mutant pancreatic tumors. Nature Medicine, 23(1):60–68, 2017.

43. Minjie Wang, D. Zheng, Zihao Ye, Quan Gan, Mufei Li, Xiang Song, Jinjing Zhou, Chao Ma, Lingfan Yu, Yu Gai, et al. Deep graph library: A graph-centric, highly-performant package for graph neural networks. arXiv preprint 1909.01315, 2019.

44. Ruiping Wang, Yue Han, Zhangxiang Zhao, Fan Yang, Tingting Chen, Wenbin Zhou, Xianlong Wang, Lishuang Qi, Wenyuan Zhao, Zheng Guo, et al. Link synthetic lethality to drug sensitivity of cancer cells. Briefings in Bioinformatics, 20(4):1295–1307, 2019.

45. Shike Wang, Fan Xu, Yunyang Li, Jie Wang, Ke Zhang, Yong Liu, Min Wu, and Jie Zheng. KG4SL: knowledge graph neural network for synthetic lethality prediction in human cancers. Bioinformatics, 37(Supplement 1):i418–i425, 2021.

46. Tim Wang, Haiyan Yu, Nicholas W Hughes, Bingxu Liu, Arek Kendirli, Klara Klein, Walter W Chen, Eric S Lander, and David M Sabatini. Gene essentiality profiling reveals gene networks and synthetic lethal interactions with oncogenic Ras. Cell, 168(5):890–903, 2017.

47. David S Wishart, Yannick D Feunang, An C Guo, Elvis J Lo, Ana Marcu, Jason R Grant, Tanvir Sajed, Daniel Johnson, Carin Li, Zinat Sayeeda, et al. DrugBank 5.0: a major update to the DrugBank database for 2018. Nucleic Acids Research, 46(D1):D1074–D1082, 2018.

48. Alan SL Wong, Gigi CG Choi, Cheryl H Cui, Gabriela Pregernig, Pamela Milani, Miriam Adam, Samuel D Perli, Samuel W Kazer, Aleth Gaillard, Mario Hermann, et al. Multiplexed barcoded CRISPR-Cas9 screening enabled by CombiGEM. Proceedings of the National Academy of Sciences, 113(9):2544–2549, 2016.

49. Hao Ye, Xiuhua Zhang, Yunqin Chen, Qi Liu, and Jia Wei. Ranking novel cancer driving synthetic lethal gene pairs using TCGA data. Oncotarget, 7(34):55352, 2016.

50. Mahdi Zamanighomi, Sidharth S Jain, Takahiro Ito, Debjani Pal, Timothy P Daley, and William R Sellers. GEMINI: a variational Bayesian approach to identify genetic interactions from combinatorial CRISPR screens. Genome Biology, 20(1):137, 2019.

51. Biyu Zhang, Chen Tang, Yanli Yao, Xiaohan Chen, Chi Zhou, Zhiting Wei, Feiyang Xing, Lan Chen, Xiang Cai, Zhiyuan Zhang, et al. The tumor therapy landscape of synthetic lethality. Nature Communications, 12(1):1–11, 2021.

52. Dongxin Zhao, Mehmet G Badur, Jens Luebeck, Jose H Magaña, Amanda Birmingham, Roman Sasik, Christopher S Ahn, Trey Ideker, Christian M Metallo, and Prashant Mali. Combinatorial CRISPR-Cas9 metabolic screens reveal critical redox control points dependent on the KEAP1-NRF2 regulatory axis. Molecular Cell, 69(4):699–708, 2018.

